# A spatiotemporally resolved single cell atlas of the *Plasmodium* liver stage

**DOI:** 10.1101/2021.12.03.471111

**Authors:** Amichay Afriat, Vanessa Zuzarte-Luís, Keren Bahar Halpern, Lisa Buchauer, Sofia Marques, Aparajita Lahree, Ido Amit, Maria M. Mota, Shalev Itzkovitz

## Abstract

Malaria infection involves an obligatory, yet clinically silent liver stage^1,2^. Hepatocytes operate in repeating units termed lobules, exhibiting heterogeneous gene expression patterns along the lobule axis^3^, but the effects of hepatocyte zonation on parasite development have not been molecularly explored. Here, we combine single-cell RNA sequencing^4^ and single-molecule transcript imaging^5^ to characterize the host’s and parasite’s temporal expression programs in a zonally-controlled manner for the rodent malaria parasite *Plasmodium berghei* ANKA. We identify differences in parasite gene expression in distinct zones, and a sub-population of periportally-biased hepatocytes that harbor abortive infections associated with parasitophorous vacuole breakdown. These ‘abortive hepatocytes’ up-regulate immune recruitment and key signaling programs. They exhibit reduced levels of *Plasmodium* transcripts, perturbed parasite mRNA localization, and may give rise to progressively lower abundance of periportal infections. Our study provides a resource for understanding the liver stage of *Plasmodium* infection at high spatial resolution and highlights heterogeneous behavior of both the parasite and the host hepatocyte.

## Main

Malaria is a mosquito-borne disease caused by *Plasmodium* spp. parasites. When an infected female *Anopheles* mosquito bites a mammalian host, it inoculates around 100 sporozoites, a motile form of the parasite capable of target recognition and host-interaction^1^. The sporozoite travels in the bloodstream until reaching the liver^2^. Upon colonizing a hepatocyte, the parasite encapsulates itself in the host’s plasma membrane forming a parasitophorous vacuole^6^, where it rapidly replicates to form a coenocyte of a few thousand nuclei^1^. At the end of the liver stage of the parasite’s life cycle (48-60 hours post-infection, hpi^7^), thousands of individual merozoites are released into the blood-stream, giving rise to the pathological blood stage.

The mammalian liver exhibits spatial heterogeneity. It is composed of repeating anatomical units termed lobules. Each lobule has a diameter of around half a millimeter in mice and consists of 9-12 concentric layers of hepatocytes^3^. Blood flows from portal nodes through radial sinusoidal channels into draining central veins, creating gradients of oxygen, nutrients and hormones. As a result, hepatocytes at different zones exhibit different gene expression signatures^8^. Periportal hepatocytes engage in protein secretion, ureagenesis and gluconeogenesis, whereas pericentral hepatocytes specialize in processes such as bile acid production, xenobiotic metabolism and glutamine biosynthesis^3^. Previous *ex-vivo* studies suggested that the pace of *Plasmodium* infection could differ between pericentral and periportal hepatocytes^9,10^. Studies using bulk RNA measurements characterized the transcriptomes of host and parasite during the liver stage of infection^11,12^, and a malaria single cell atlas was generated using an *ex-vivo* platform^13^. However, accounting for the hepatocyte spatial heterogeneity and identifying heterogeneous host and parasite responses requires *in-vivo* single cell approaches.

### A single cell atlas of the Plasmodium liver stage

To study the *Plasmodium* liver stage at single cell resolution we injected mice with GFP-expressing *Plasmodium P. berghei* ANKA^14^ (Fig. 1a). We sacrificed infected mice at different time points post-infection (2, 12, 24, 30 and 36hpi), and extracted livers for single cell RNA sequencing (scRNAseq, Extended Data Fig. 1). We also embedded liver tissue for single molecule fluorescence *in-situ* hybridization (smFISH) experiments. We sorted GFP+ and GFP-hepatocytes and performed scRNAseq using the MARS-seq protocol^4^. We aligned the reads to both mouse and *Plasmodium* genomes, obtaining 13,457 hepatocytes of which 2,424 were infected (Fig. 1b,c). Both infected and uninfected hepatocytes exhibited clear zonated expression programs, as evident from the mutually exclusive expression of the periportally-zonated hepatocyte gene Cyp2f2 and the pericentrally-zonated gene Cyp2e1 (Fig. 1d). We used a previously established set of hepatocyte landmark genes^15^, and additionally filtered for those that did not exhibit changes between infected and uninfected hepatocytes to establish a zonation score for each hepatocyte that is correlated with its location along the lobule radial axis (Methods, Fig. 1e). At 2hpi, both infected and uninfected hepatocytes exhibited a global elevation of genes associated with the stress involved in tissue dissociation^16^, including genes that we validated not to increase post-infection *in-situ* (Extended Data Fig. 2). We therefore excluded this early time point from further analyses of hepatocyte response to infection.

**Fig. 1:**
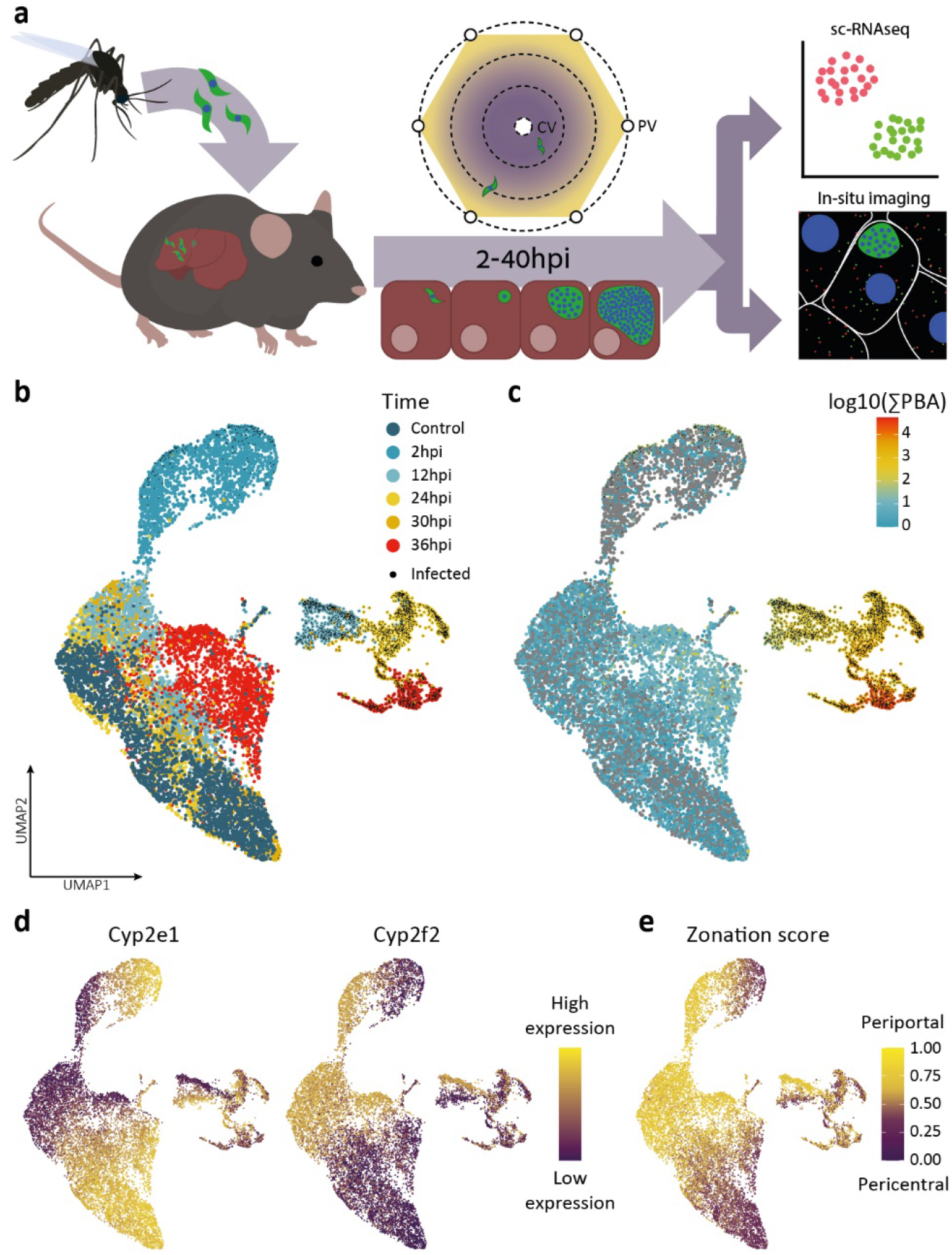
A single cell atlas of the *Plasmodium* liver stage enables annotation of infections by zone and time. **a**, GFP+ parasites are injected into mice, livers are extracted at different ensuing time points for scRNAseq and in-situ imaging. CV/PV - central/portal vein. **b**, UMAP of hepatocytes colored by hours post-infection (hpi). **c**, UMAP of hepatocytes colored by log10 of the sum of *Plasmodium* (PBA) reads. Black dots in (c-d) denote infected hepatocytes. **d**, UMAP colored by the expression of the pericentral hepatocyte gene Cyp2e1 and the periportal gene Cyp2f2. **e**, Zonation score inferred from the sum of zonated hepatocyte landmark genes. UMAP projections reconstructed based on the combined mouse and *Plasmodium* transcriptomes.

### Hepatocyte response to malaria infection

To identify host programs modified by infection at different time points, we performed differential gene expression analysis between the infected and uninfected hepatocytes. Importantly, we stratified single hepatocytes by their inferred zone so that comparisons were performed between infected and uninfected hepatocytes that reside at similar lobule coordinates and were sampled at the same time points (Methods, Fig. 2a, Extended Data Fig. 3, Supplementary Table S1). We found that infected hepatocytes up-regulated genes enriched for several programs including TNFa signaling, interferon alpha and gamma responses and glutathione metabolism (Fig. 2b). Up-regulated genes included Ftl1 and Fth1, encoding the iron chelator ferritin light and heavy chains and Slc40a1, encoding the iron export ferroportin-1 transporter (Fig. 2a). Iron is essential for liver-stage malaria development, and the iron chelator ferritin has been shown to increase in the serum of infected individuals^17^. The elevated levels of hepatocyte ferritin and ferroportin genes might be an adaptation to deprive the parasite of available iron. Down-regulated processes in infected hepatocytes included fatty acid metabolism, bile acid metabolism and complement and coagulation cascades (Fig. 2b). Given the essential role of fatty acids for the parasite development^18,19^, the reduction in hepatocyte fatty acid biosynthesis genes such as Acly and Fasn (Fig. 2a) might serve to deprive it from these key building blocks. We used smFISH to validate the predicted change in expression of representative genes in infected hepatocytes (Fig. 2c-f), demonstrating a significant increase in Ftl1 and a decrease in G6pc, Fasn, and Apob.

**Fig. 2:**
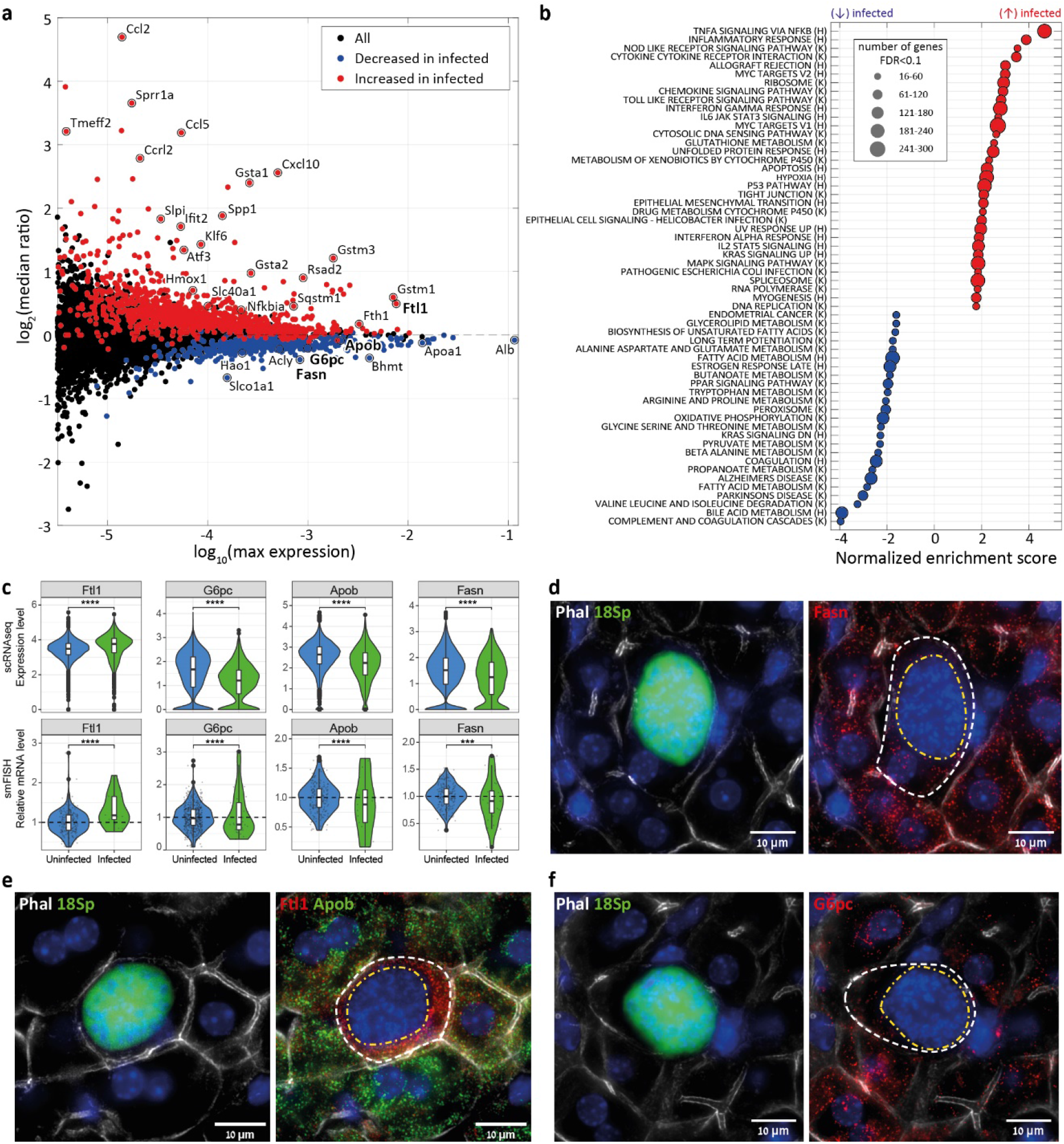
Zonally-stratified comparisons identify up-regulated genes in infected hepatocytes. **a**, MA plot showing the median expression ratio over time and space between infected and uninfected hepatocytes. Cells were binned by time (12/24/30+36 hpi) and zone (PC or PP), ratios shown for each gene are medians of the ratios over all time points. Y axis indicates log2 of the median ratio per gene. X axis indicates log10 of the gene’s max expression across all time and zone bins. Genes significantly increased or decreased in infected hepatocytes are plotted in red or blue respectively (FDR q-value<0.01, Methods). **b**, Gene set enrichment analysis (GSEA) shows increase in immune and stress pathways alongside a decrease in multiple metabolic pathways in infected hepatocytes. (H) denote Hallmark gene sets, (K) denote KEGG gene sets. **c**, Quantification of smFISH experiments validate predicted differentially expressed hepatocyte genes. Significance was determined by one-sided Wilcoxon rank-sum test (**:p<=0.01; ***:p<=0.001; ****:p<=0.0001, Methods). Boxes outline the 25-75 percentiles, horizontal black lines denote the median. **d-f**, smFISH images of validated genes at 40hpi. Left panel - Phalloidin (Phal) in white, parasite Ch12 18S rRNA (18Sp, PBANKA_1245821) in green, Dapi blue; Right panel - Fasn/Ftl1/G6pc mRNA in red, Apob mRNA in green. White dashed line indicates infected hepatocyte; Yellow dashed line indicates parasite.

To identify potential zonated patterns in infection rates we analyzed the computationally-inferred zonation scores of infected hepatocytes (Extended Data Fig. 4a). The single cell data suggested that pericentral infections were more abundant at all-time points, however this differential abundance could stem from differences in the efficiency of single cell extraction from different lobule zones. To identify zonated features of infection rates in an unbiased manner, we therefore analyzed the zonal abundances of infected hepatocytes *in-situ*. We combined smFISH for the periportally-zonated albumin-encoding gene Alb^8^, and established an *in-situ* zonation score based on Alb levels of hepatocytes that neighbor each infected cell (Extended Data Fig. 4b-d, Methods). We found that infected hepatocytes were not zonated at 2, 15, 24 and 36hpi but were significantly more abundant in the pericentral zones at 40hpi (Extended Data Fig. 4d). We further used our scRNAseq data to demonstrate that the parasite mRNA content was significantly higher in pericentral infections specifically at the latest sequenced time point of 36hpi (Extended Data Fig. 4e). Parasites infecting pericentral hepatocytes therefore seem to survive and to develop at a higher rate compared to parasites in periportal hepatocytes.

### Abortive hepatocytes exhibit a distinct molecular signature

What could be the reason for the lower abundance and smaller fraction of parasite transcripts in periportal hepatocytes at late time points? To address this question, we examined the scRNAseq data of infected hepatocytes at 36hpi (Fig. 3a). We found that infected hepatocytes exhibited two distinct clusters based on the host transcriptome. The minor cluster was enriched in periportal hepatocytes (Fig. 3b) and contained cells with significantly lower numbers of parasite reads (Fig. 3c). This minor cluster was enriched in genes related to immune programs (Fig. 3d-f), such as Cxcl10, Nfkbia and Sqstm1, P53-pathway genes such as Mdm2, the transcription factor Myc and its downstream targets, and the Notch downstream target transcription factor Hes1 (Fig. 3d,e-h).

**Fig. 3:**
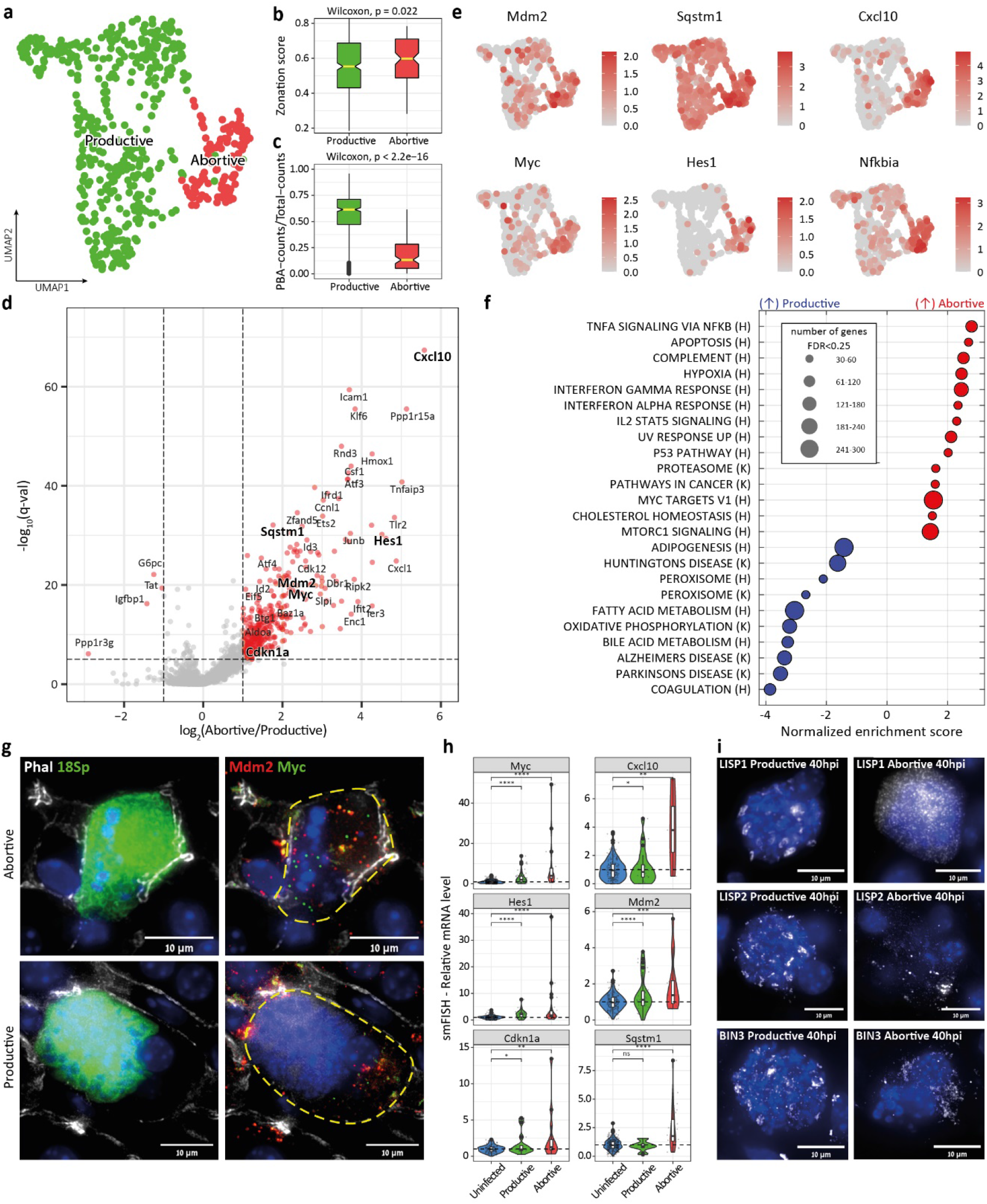
A periportally-enriched sub-population of infected abortive hepatocytes is associated with vacuole breakdown. **a**, Infected hepatocytes exhibit a distinct sub-cluster at 36hpi, annotated as ‘abortive hepatocytes’. **b**, Abortive hepatocytes are more periportally zonated compared to productive hepatocytes. Shown are the computationally-inferred zonation scores. **c**, Abortive hepatocytes harbor a smaller fraction of parasite mRNA compared to productive hepatocytes at 36hpi. For both (**b**) and (**c**) boxes outline the 25-75 percentiles, horizontal yellow lines denote the median. **d**, Volcano plot showing differentially-expressed hepatocyte genes between the two clusters. Bold genes are highlighted in (**e**). **e**, UMAP plots of representative genes up-regulated in abortive hepatocytes. **f**, GSEA analysis demonstrates elevation of immune-recruiting pathways, Myc and P53 pathways. **g**, Abortive cells harbor a disintegrated vacuole. Left panel - Phalloidin (Phal) in white, parasite Ch12 18S rRNA (18Sp, PBANKA_1245821) in green, Dapi blue; Right panel – Mdm2 mRNA in red, Myc mRNA in green. **h**, Quantification of smFISH images validating the increase in the abortive cluster host genes. Significance was determined by one-sided Wilcoxon rank-sum test (ns:p>0.05; *:p<=0.05; **:p<=0.01; ***:p<=0.001; ****:p<=0.0001, Methods). Boxes outline the 25-75 percentiles, horizontal black lines denote the median. **i**, Abortive parasites exhibit loss of mRNA localization for some *Plasmodium* genes. UMAP projections in (**a**) and (**e**) reconstructed based on the mouse transcriptomes of the infected hepatocytes at 36hpi.

Our *in-situ* validations of the signatures of the minor cluster (Fig. 3g,h), identified by their elevated levels of marker transcripts (Fig. 3d-e, Extended Data Fig. 5a-d), uncovered distinct morphological features of the minor cluster cells. The parasitophorous vacuole was disintegrated, as evident by the ubiquitous fluorescence of both GFP and parasite 18S rRNA (Fig. 3g). The parasitic nuclei were also dispersed throughout the hepatocyte cytoplasm. Our in-situ analysis validated the portal enrichment of these cells (Extended Data Fig. 5e). The fractions of hepatocytes with disintegrated vacuole increased from 3% at 24hpi through 17% at 36hpi and 27% at 40hpi (Extended Data Fig. 5e). These events most likely do not represent hepatocytes harboring productive merozoites, since merozoite formation and release from hepatocytes starts at 50-60hpi in-vitro^7^. Moreover, blood transfer from liver-stage mice gave rise to parasitaemia in recipient only at 52hpi and not in 42hpi^20^. Given the pattern of vacuole breakdown and relatively early phase of their appearance, we therefore termed these cells ‘abortive hepatocytes’.

Abortive hepatocytes harbored a distinct parasite gene expression signature compared to productive hepatocytes (Supplementary Table S2) that included higher expression of the *Plasmodium* heat shock proteins HSP90, HOP and UIS24 (Extended Data Fig. 5f). We also identified distinctly different mRNA localization patterns for several parasite genes between abortive and non-abortive hepatocytes - the *Plasmodium* transcripts for LISP1, LISP2 and BIN3 (PBANKA_090330) were localized in distinct foci in productive hepatocytes, yet were completely interspersed in abortive hepatocytes (Fig. 3i). Our analysis therefore highlights a molecular blueprint of periportally-biased infected hepatocytes with an abortive phenotype and elevated expression of immune-recruiting programs. Elimination of these abortive hepatocytes by the immune system could account for the lower abundance of periportally-infected hepatocytes at late time points of infection.

### Spatio-temporal expression programs of the parasite

We next used our single cell atlas to examine the developmental programs of the parasite during the liver stage (Fig. 4a,b, Extended Data Fig. 6, Supplementary Table S3). We found that parasite genes clustered into distinct temporal profiles, enriched for specific gene sets. Early parasite genes included RNA polymerases (Extended Data Fig. 6c) and purine and pyrimidine metabolism genes (Fig. 4c), crucial for the synthesis of mRNA during the ensuing massive parasite proliferation. Early genes also included ubiquitin genes (Fig. 4c), presumably serving to erase the protein content of the preceding sporozoite state (Extended Data Fig. 6a). The parasites next sequentially up-regulated DNA polymerases (Extended Data Fig. 6d) and metabolic programs for glycolysis, TCA cycle, biotin and glutathione metabolism (Fig. 4e,f, Extended Data Fig. 6e-g). Lastly, parasites induced genes associated with fatty acid metabolism and apicoplast-related pathways such as chlorophyll metabolism (Fig. 4g, Extended Data Fig. 6h). The late increase in the parasite genes encoding de-novo fatty acid biosynthesis coincided with a decline in the earlier expression of the genes encoding UIS4 and UIS3 (Fig. 4b, Extended Data Fig. 6a), which have been suggested to facilitate transport of free fatty acids from the hepatocyte host^21^. We used smFISH to validate the temporal programs of selected parasite genes (Fig. 4h-j).

**Fig. 4:**
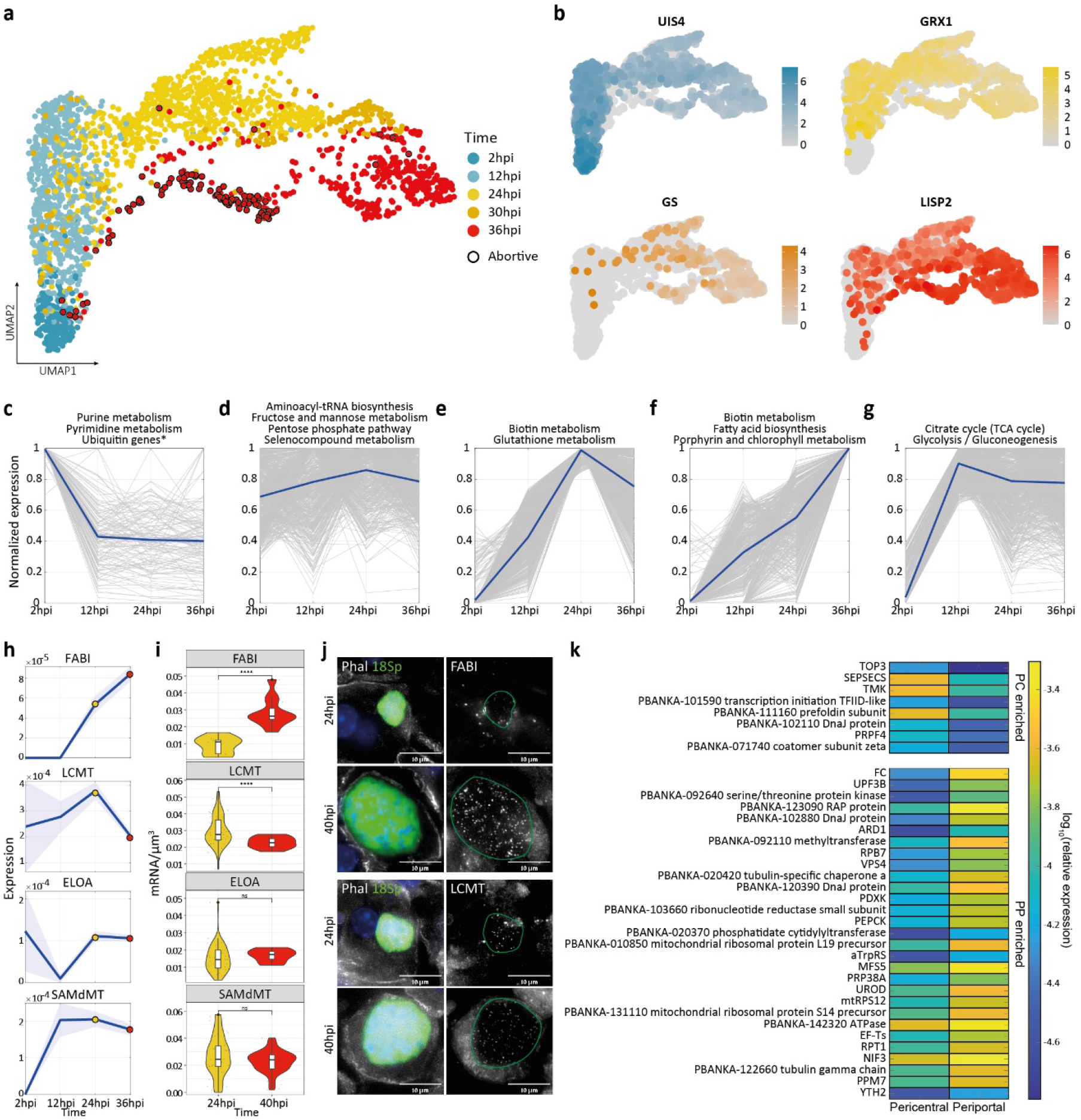
Temporally-resolved programs of the *Plasmodium* transcriptome. **a**, UMAP projections of the infected hepatocytes computed based on the *Plasmodium* transcriptomes, colored by hours post-infection. Abortive cells annotated according to the minor hepatocyte cluster in Fig 3a. **b**, Representative genes color coded by the stage of infection in which expression peaks – early gene (UIS4, blue), intermediate early gene (GRX1, yellow), intermediate late gene (GS, orange) and late gene (LISP2, red). **c-g**, K-means clustering of *Plasmodium* averaged expression programs. Titles are *Plasmodium* KEGG pathways enriched in each temporal cluster. An exception is the early Ubiquitin genes set, which was the major component of the enriched Arginine biosynthesis and Tryptophan biosynthesis pathways (Supplementary Table S3). Grey– temporal profile of genes in cluster, Blue – mean temporal expression over all genes in cluster. Expression of all genes in **c**-**g** normalized to the maximum across time points. **h**, Temporal expression profiles of *Plasmodium* genes used for smFISH validation. Light patches denote the SEM. Yellow and red dots indicate validated time points in **i-j**, smFISH quantification validates temporal prediction. (**i**) Significance was determined by two-sided Wilcoxon rank-sum test (ns:p>0.05; ****:p<=0.0001, Methods). Boxes outline the 25-75 percentiles, horizontal black lines denote the median. (**j**) Left panel - Phalloidin (Phal) in white, Ch12 18S rRNA (18Sp, PBANKA_1245821) in green, Dapi blue; Right panel – FABI (PBANKA_122980)/LCMT(PBANKA_130360 leucine carboxyl methyltransferase) mRNA in white, green line indicates vacuole outline. **k**, Zonally enriched *Plasmodium* genes at 36hpi (FDR q-val<0.3, max relative expression>5×10^−5^).

We further identified *Plasmodium* genes that were differentially expressed between pericentral and periportal hepatocyte host cells (Supplementary Table S2, Fig. 4k). Parasites in periportal hepatocytes exhibited higher levels of genes involved in heme biosynthesis, including Uroporphyrinogen decarboxylase (UROD)^22^ and FC, encoding ferrochelatase^23^. Heme is essential for liver-stage *Plasmodium* development. Hepatocyte heme biosynthesis is pericentrally-zonated^8,24^, as it is part of the pericentrally-abundant cytochrome-P450. Periportal hepatocytes might therefore offer less available heme for salvation by the parasite, necessitating its de-novo synthesis.

### Discussion

Our study provides a comprehensive overview of the life cycle of individual parasites throughout the *Plasmodium* liver stage. The lack of zonated abundances at early time points indicate that the parasite does not seem to preferentially colonize hepatocytes at specific zones. Rather, our results align with a random process of colonization, yet zone-dependent rates of development and/or survival. The higher pericentral abundance of infections at late time points could be explained either by the lower oxygen tension at the pericentral lobule layers^3^, which has been shown to promote parasite development *ex-vivo*^9^, or due to lower survival of periportal infected hepatocytes. The liver lobule exhibits immunological zonation^25^, with a higher periportal abundance of myeloid and lymphoid cells. The decreased frequencies of periportal infections could be explained by higher elimination rates of abortive hepatocytes, which we found to be more periportally abundant. Indeed, we found that abortive hepatocytes exhibit a distinct gene expression signature enriched in immune recruitment genes, such as interferon-gamma response and to a lesser extent, interferon-alpha response. Type I interferon has been shown to mediate liver-stage immune response^20,26^, as well as contribute to immune mediated pathology^27^. The abortive cells we have identified may eventually become immune-infiltrated and eliminated^20^. It will be interesting to apply paired-cell approaches^15,28^ to explore the interactions between abortive hepatocytes and specific immune cell subsets.

Our scRNAseq data suggested that abortive hepatocytes strongly elevate key pathways, such as Notch, P53 and Myc. P53 inhibition has been shown to affect liver stage malaria progression^29^, and Myc inhibition dampens acute liver failure^30^. It would be interesting to apply our approach of spatially resolved scRNAseq of infected hepatocytes in mice with drug-induced or genetically induced perturbations of these pathways. We have identified several potentially adaptive programs of the host hepatocyte and the parasite, including processes such as heme, iron and fatty acid metabolism, and zonal trends in both the host and the parasite. The combined measurements of the parasite and host forms a resource for the detailed analysis of the *Plasmodium* liver stage, while accounting for the liver’s spatial heterogeneity^3^. It can serve as a basis for exploration of potential vulnerabilities and identification of targetable host and parasite pathways^31^.

## Methods

### Mice and tissues

Experiments were conducted on 6-7 week old C57BL/6J female mice. Mice were purchased from the Charles River Breeding Laboratories and were housed in the facilities of the Instituto de Medicina Molecular in Lisbon in a germ-free environment supplied with water and food ad libitum. All *in vivo* protocols were approved by the internal animal care committee of Instituto de Medicina Molecular and were performed according to national and European regulations. For smFISH, 2 mice were sampled per time point. For scRNAseq 4/2/3/1/2 mice were sampled for 2hpi/12hpi/24hpi/30hpi/36hpi respectively, with 2 mice for control (one none infected and one mock infected). Due to technical limitations, mice from time points 24hpi and 12hpi were collected ±1-2 hours of the designated time point.

### Parasite

GFP expressing *Plasmodium berghei ANKA* (clone 259cl2^14^) were used. Sporozoites were obtained through dissection of the salivary glands of infected female Anopheles stephensi mosquitoes bred and infected at the IMM. Mice were inoculated using retro-orbital injection. For scRNAseq, each mouse was injected with 10^6^ sporozoites in 200µl DMEM. Mock infected control mouse was injected with filtered mosquito salivary gland debris devoid of the parasite. For smFISH, each mouse was injected with 2×10^4^ sporozoites in 200µl DMEM.

### Liver dissociation and FACS sorting of hepatocytes

Mice livers were perfused and dissociated into single cells using Liberase Blendzyme 3 recombinant collagenase (Roche Diagnostics) as previously described^8^. Isolated hepatocytes were sorted on BD FACS Aria IIu using a 130 μm nozzle and 1.5 neutral density (ND) filter. Samples were stained with αCD45, αCD31 antibodies and DAPI or PI. Cells were gated to include live cells only (DAPI/PI negative) and exclude doublets (FSC, SSC) and non-parenchymal (CD31- and CD45-). The remaining cells were then gated for infected (GFP+) or uninfected (GFP-) and sorted accordingly. The cells were sorted into 384-well capture plates containing 2μl lysis solution and barcoded poly(T) reverse-transcription (RT) primers for MARS-seq^4^, allowing for both single cell barcoding and unique molecular identifiers (UMIs) barcoding of mRNA transcripts. Every plate contained uninfected hepatocytes and several rows (3,5 or 10) of infected hepatocytes. Four wells were left empty on the bottom left corner of each plate (wells O1, O2, P1 and P2) for background control. Sorted plates were spun down, frozen on dry ice and then kept at - 80°C until library preparation.

### MARS-seq library preparation and sequencing

Libraries were prepared as previously described^4^. Briefly, mRNA in capture plates were barcoded and reverse transcribed into cDNA then pooled together. The pooled cDNA libraries were then amplified using T7 in-vitro transcription and fragmented. Resulting RNA libraries were then tagged with pool-specific barcodes and illumina sequencing adapters and converted to cDNA again. Pooled libraries were quality controlled at different times of the protocol and prior to sequencing. The final libraries were then pooled together (15-20 at a time) and sequenced using the NextSeq 500/550 kit High Output Kit v2.5 (Illumina 20024906). The Illumina output files were converted to fastq format using bcl2fastq (v2.20.0.422) and then aligned to a combined reference genomes of *Mus musculus* (GRCm38.p6) and *Plasmodium berghei ANKA* (PBANKA01.43) using STAR (v2.7.3a) and zUMI (v0.0.6c).

### scRNAseq data processing

Data processing was done on Python (3.7.6). For every plate, the background counts per gene were calculated based on the mean expression in the empty wells (defined as wells with less than 1,000 total UMIs). The background was then subtracted from all the wells in the plate. Using the Ensembl database, rows were renamed using ‘gene-id’ indication and pseudogenes were filtered out based on the ‘gene-biotype’ indication. Rows with duplicated gene-ids were merged into a single row. Cells were divided into ‘infected’ and ‘uninfected’ based on sorting scheme per plate. Since sorting/barcoding errors occasionally led to miss-classification of the infected wells, a classifier was further applied based on the logarithm of the ratio between the summed PBA UMIs and the total cellular UMIs (after adding a pseudocount of 1 to both). For each plate, 100 thresholds for this ratio were scanned, and for each threshold, false positive rates (FPR) and false negative rates (FNR) were computed. FPR was defined as the fraction of cells annotated as uninfected that were above the threshold. FNR was defined as the fraction of cells annotated as infected that were below the threshold. The equal error rate was selected as the threshold for which FPR=FNR. Only cells annotated as infected/uninfected that had a log PBA summed ratio above/below this threshold respectively were retained.

Following the filtered annotation, a PBA transcriptomic background was subtracted from all wells in plates that included infected cells. To this end, the annotated uninfected cells were considered as the background. The 99^th^ percentile of every PBA gene in uninfected cells was subtracted from the same gene in all cells. The pooled and background-subtracted dataset was next filtered for high mitochondrial content. The fraction of mouse mitochondrial genes was calculated for every remaining cell, and cells with mitochondrial fraction above the 95 percentile per mouse were removed. Cells were next filtered to retain cells with more than 1,500 and less than 150,000 reads, as well as more than 200 and less than 8,000 genes. Based on the filtered cells, low yielding plates were excluded as follows – For each plate the ratio between the median log summed UMI counts per plate and the median log summed UMI counts per mouse was computed. A Z-score was then calculated for this value, and plates with Z-score of <-1.5 were removed from the dataset. From this point onward, all analyses were performed on the reads of protein-coding genes only, excluding mouse mitochondrial genes and Major urinary proteins (Mups) known to be highly variable between mice^32^. In addition, remaining ‘infected’ cells with less than 30 PBA-reads were filtered out.

### scRNAseq processed data analysis

The processed UMI table was analyzed using Seurat 4.0.1^33^ running on R4.0.2. Data normalization and scaling followed the suggested default settings for most of the package functions. The data were log-normalized and scaled without regression. Top 2,000 variable genes were selected using “FindVariableFeatures” function with the “vst” method. Principle component analysis was based on these genes, and the first 10 components were used for clustering (0.4 resolution) and dimension reduction (UMAP, using the ‘cosine’ metric).

Subsets of the main Seurat structure were used to cluster and analyze the 36hpi infected cells using only mouse transcripts (Fig. 3), and all the infected cells using PBA transcripts (Fig. 4). The subsetted cells were renormalized based on the respective subset of transcripts. For each subset, the top 2,000 variable genes were selected, and the first 10 principle components were used for clustering (resolution 0.1 in Fig. 3 and 0.4 in Fig. 4) and UMAP reductions were generated (using ‘cosine’ metric in Fig. 3 and ‘euclidian’ metric in Fig. 4). For differential gene expression analysis and pathway enrichment, the raw data was normalized to relative counts per cell in MATLAB R2019a by dividing by the sum of all genes that individually take up less than 0.02 of the cellular summed UMIs when averaged over all cells.

### Zonation reconstruction

Single-cell spatial locations along the lobule axis were inferred computationally from the data based on the sum of a panel of landmark genes, as previously described^15^. However, for this study a smaller subset of landmark genes was used, retaining only genes that showed no significance change in expression between infected and unifected cells. Periportal landmark genes used were: Apof, Apom, Asgr2, Atp5a1, C1s1, C8b, Cpt2, Tkfc, Eef1b2, Fads1, Gc, Hsd17b13, Ifitm3, Igf1, Igfals, Ndufb10, Pigr, S100a1, Serpina1c, Serpina1e, Serpind1, Serpinf1, Uqcrh, Vtn, Arg1, Cps1. Pericentral landmark genes used were: Alad, Aldh1a1, Nat8f2, Cpox, Cyb5a, Cyp3a11, Lect2, Mgst1, Prodh, Slc16a10.

### Stratified ratio change - MA-plot

To find global markers of Liver-stage infection while excluding spatio-temporal biases, cells were binned based on their metadata into 3 time points (12/24/30+36hpi) and 2 zones (pericentral/periportal = zonation score lower/higher than the 30^th^ percentile of overall zonation scores). The ratio change of mean normalized expression between infected and uninfected cells was calculated for every mouse gene (Supplementary Table S1) and one-sided Wilcoxon rank-sum tests were used to establish statistical significance of elevated or reduced expression in infected cells. Global ratio change was calculated as the median value of the ratio-changes per gene in each of the 6 bins. Overall significance was calculated using Fisher combined method corrected using Benjamini-Hochberg FDR<0.01.

### Pathways enrichment

For mouse gene set enrichment, genes with mean relative expression > 10^−5^ of summed mouse UMIs were ranked based on their ratio change between cell subsets (infected/uninfected; abortive/productive; pericentral/periportal; etc.). The ranked ratio was the basis for Gene Set Enrichment Analysis (GSEA, v3.0)^34^. Curated KEGG and HALLMARK annotations were used, filtered for minimum 15 genes in set and maximum 500. Default setting of 1,000 permutation was used to establish significance. For abortive cells DGE stricter thresholds were used - relative expression > 10^−4^, and minimum 30 genes in set.

KEGGREST (v1.28.0) was used in R4.0.2 to assign PBA genes to different Malaria KEGG pathways. Pathways were grouped into 5 clusters based on their peak expression time-point using k-means clustering in MATLAB (‘Distance’ metric = ‘cosine’).

### smFISH

Mice were sacrificed by cervical dislocation and their livers harvested. Tissues were fixed in 4% paraformaldehyde (Santa Cruz Biotechnology sc-281692) for 3 h, incubated overnight with 30% sucrose in 4% paraformaldehyde, embedded in OCT (Tissue-Tek #4583) and stored at -80°C. 8-15µm thick cryosections were used for probe hybridization as previously described^35^. Briefly, the sections were permeabilized in cold 70% ethanol for 2 hours, then rehydrated in 2XSSC (Ambion AM9763). Rehydrated tissues were treated with proteinase K (10 µg/ml Merck 124568) and then incubated with 5% or 15% Formamide (Ambion AM9342) in 2× SSC (5% formamide concentration was used for hybridization of *Plasmodium* probe libraries with low GC content). Treated sections were then mounted with hybridization buffer (5%/15% Formamide; 10% Dextran sulfate Sigma D8906; 0.02% BSA Ambion AM2616; 1 mg/ml E.coli tRNA Sigma R1753; 2 mM Vanadyl-ribonucleoside complex NEB S1402S; 2XSSC) containing diluted probes and incubated over night at 30°C. Probe libraries (Supplementary Table S4) were coupled with Cy5 or Alexa594 and diluted to 1:3000, with the exception of the 18Sp probe that was coupled with Atto488 and was used in 1:3000/6000/30000 dilution for different stages of infection (2hpi/15hpi/24-40hpi respectively). After hybridization, the sections were incubated with 50 ng/ml DAPI (Sigma-Aldrich, D9542) in 5/15% Formamide for 30 min at 30 °C, for nuclear staining and then washed in GLOX buffer (0.01M TRIS pH 8.0 Ambion M9856; 0.4% Glucose Sigma-Aldrich G8270; 2XSSC) until mounting and imaging. Samples that required hepatocyte segmentation, underwent additional staining with 1:500 Rhodamine conjugated Phalloidin (Invitrogen R415) in GLOX for 15 minutes at RT. Imaging was performed on Nikon-Ti-E inverted fluorescence microscope using the NIS elements software AR 5.11.01. The dot stack images were first filtered with a three-dimensional Laplacian of Gaussian filter of size 15 pixels and standard deviation of 1.5 pixels. Images were used for gene expression validations or in-situ zonation analysis.

### Single-cell in-situ gene expression validations

Single field 100X images were taken of minimum 21 consecutive 0.3µm z-stacks. ImageM^35^ was used for segmentation and dot counting. For Mouse genes: Segmentation was done manually based on phalloidin staining of the hepatocyte borders. Cytoplasms of productive infected cells were segmented to exclude the parasite, while abortive infected cells were segmented as a whole. Nuclei were segmented semi-automatically by the software. Gene transcripts were counted in 10 consecutive z-stacks and divided by the total volume of the segmented cell (excluding nuclei). mRNA concentration of every infected cell was compared to those of 5-10 adjacent uninfected hepatocytes (Wilcoxon rank-sum test, one-sided) and a p-value was obtained. Fisher’s combined probability test was then used to combine these results and test the overall hypothesis per gene. For *Plasmodium* genes: the parasite vacuole was segmented based on a probe for the parasites chromosome 12 18S rRNA (18Sp, PBANKA_1245821). Gene transcripts were counted in 5-10 consecutive z-stacks and divided by the total volume of the segmented parasite. mRNA concentrations were compared between different time points using Wilcoxon rank-sum test (two-sided).

### In-situ zonation analysis

Tissue samples were hybridized with Alb mRNA Cy5 and 18Sp Atto488 probes. Scans of the whole tissue sections were imaged in 10X in a single z-stack. Single fields were autofocused and stitched together using NIS elements “Scan large image” feature. Images were manually processed to remove out-of-focus pixels and to identify the positions of infected cells. The processed images were then analyzed in MATLAB as follows: A small section of background pixels (usually central/portal vein) were marked and a background threshold established (mean pixel value+5×SDpixel-value). A 40×40 pixel window was sampled around the center of every infected cell, then a number of randomly placed 40×40 pixel windows were sampled from the image (excluding areas below the background threshold). The median pixel value of Alb mRNA signal was then calculated for every window, excluding background pixels. Then, per infected cell, the fraction of random windows with a median value lower than that of the cell was calculated and regarded as an approximate zonation score for every infected cell, 0 being very pericentral and 1 periportal. The zonation score distributions of consecutive time point were compared using two-sided Wilcoxon rank-sum tests.

## Data availability

Data generated in this study will be deposited in the Gene Expression Omnibus.

## Code availability

The code used to process the raw data to a Scanpy/Seurat structure used for analysis is available at https://github.com/AmichayAfriat/SpatioTemporal_malaria_liver_stage_atlas/. Codes for further analysis will be provided by the authors upon request.

## Acknowledgements

S.I. is supported by the Wolfson Family Charitable Trust, the Edmond de Rothschild Foundations, the Fannie Sherr Fund, the Dr. Beth Rom-Rymer Stem Cell Research Fund, the Minerva Stiftung grant, the Israel Science Foundation grant No. 1486/16, the Broad Institute-Israel Science Foundation grant No. 2615/18, the European Research Council (ERC) under the European Union’s Horizon 2020 research and innovation program grant No. 768956, the Chan Zuckerberg Initiative grant No. CZF2019-002434, the Bert L. and N. Kuggie Vallee Foundation and the Howard Hughes Medical Institute (HHMI) international research scholar award.

L.B. is supported by the European Molecular Biology Organization under EMBO Long-Term Fellowship ALTF 724-2019.

This work was also financed by “laCaixa” Foundation (HR17/52150010) to M.M.M. and Fundação para a Ciência e Tecnologia (LISBOA-01-0145-FEDER-030751 and PTDC/BIA-MOL/30112/2017) to M.M.M and V.Z.L.

## Contributions

S.I. and M.M.M. conceived the study; A.A., V.Z.L., K.B.H., S.M. and A.L. preformed the experiments; A.A., L.B. and S.I. performed the data analysis; I.A. contributed to project design; S.I., A.A. and M.M.M. wrote the paper; All of the authors discussed the results and commented on the manuscript.

## Extended Data Figures

**Extended Data Fig. 1:**
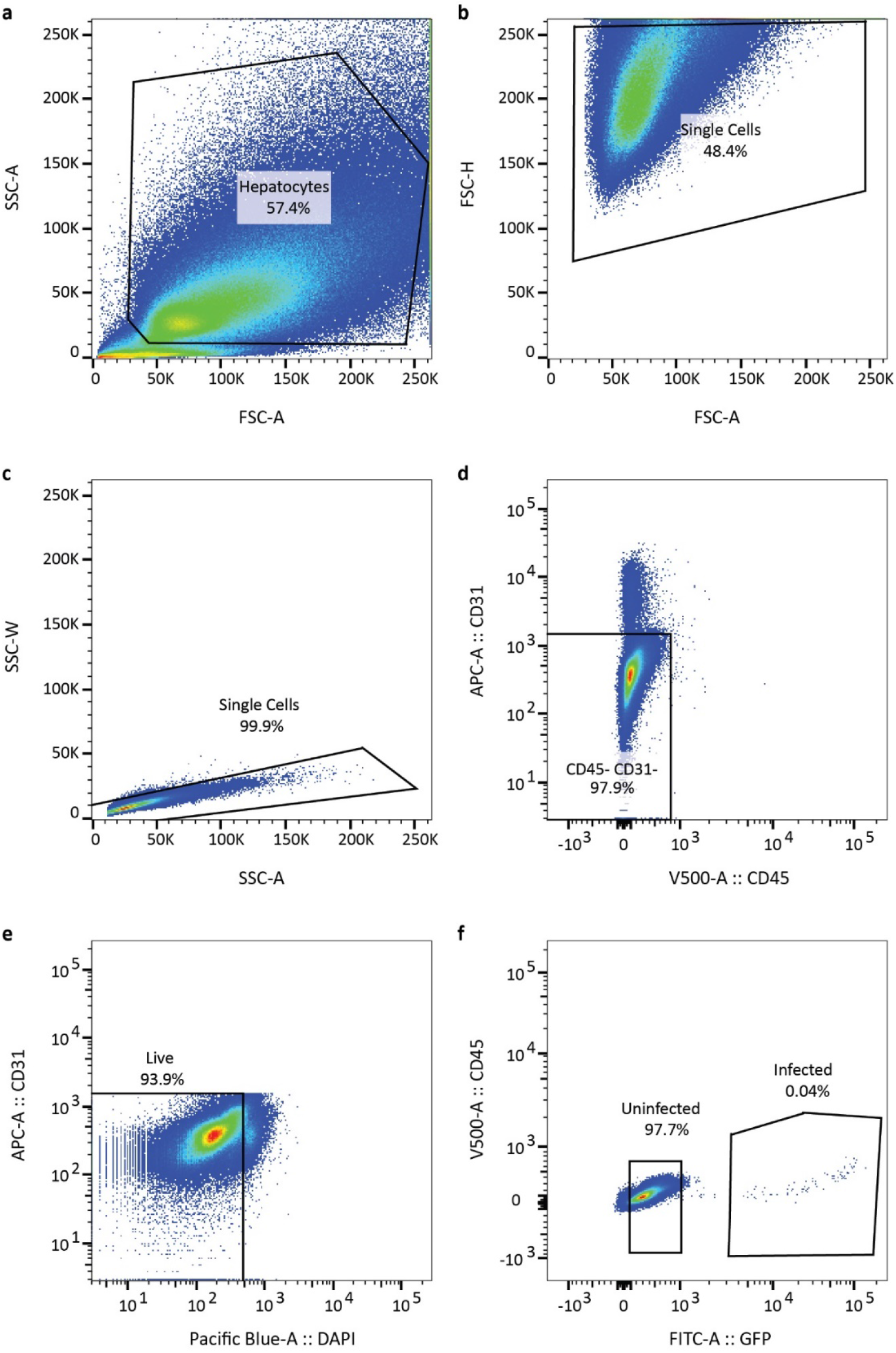
Representative FACS gating for scRNAseq. **a**, FACS gating based on general size to find hepatocytes. **b-c**, Gating to discern single-cells from clumps of cells. **d**, Negative selection to exclude non-parenchymal cells. **e**, Gating to include only live hepatocytes. **f**, gating for infected and uninfected hepatocytes based on GFP content.

**Extended Data Fig. 2:**
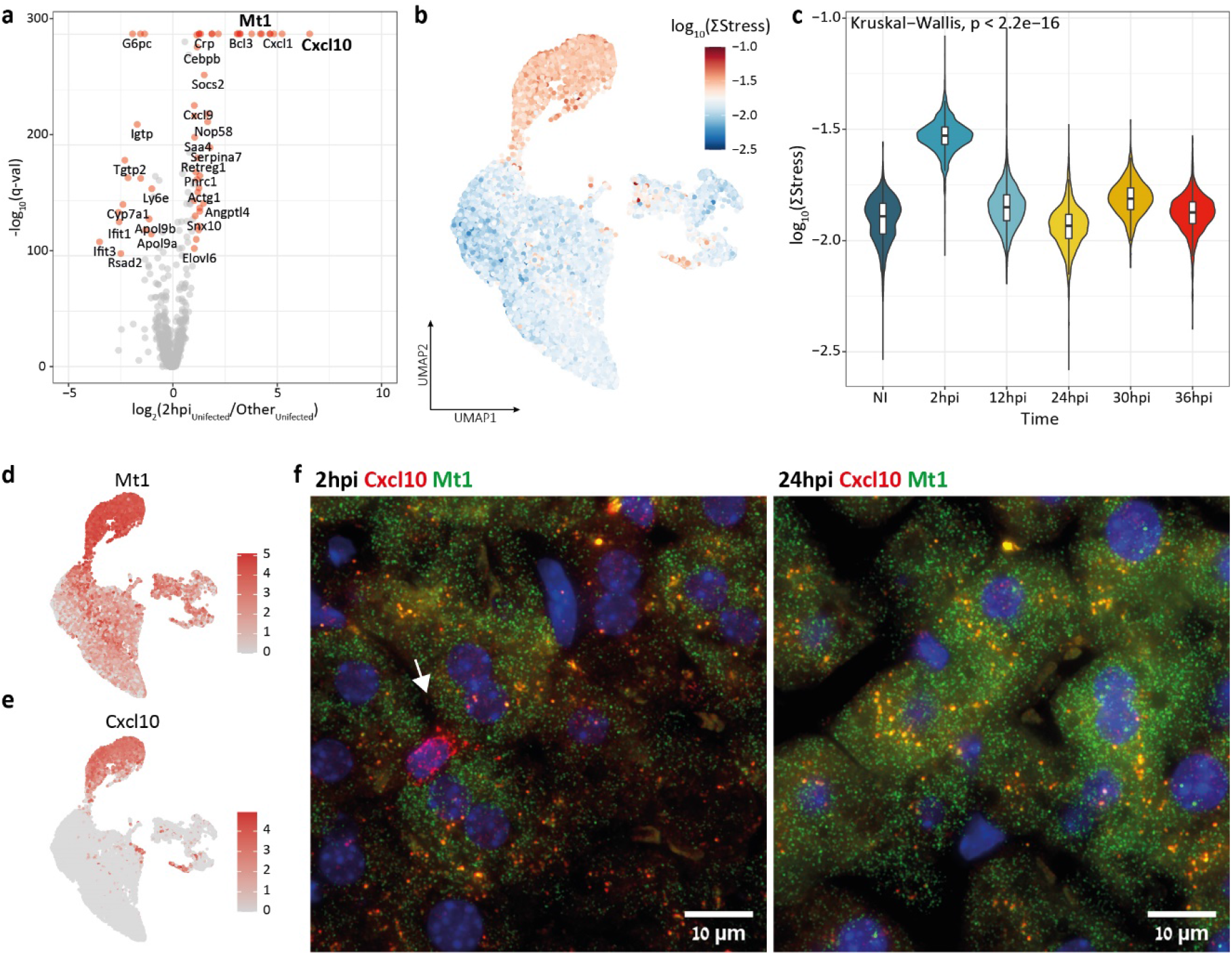
ScRNAseq of uninfected cells acquired at 2hpi exhibit an increase in genes previously shown to increase following tissue dissociation. **a**, Volcano plot showing differentially-expressed hepatocyte genes between uninfected cells acquired at 2hpi and at every other time-point. Bold genes are highlighted in (**d-f**). **b**, UMAP plot colored by log10 summed relative expression of dissociation marker genes (van den Brink et al. 2017)^16^ and (**c**) Violin plot of stress indication in uninfected cells binned by time demonstrates notable increase in dissociation stress at 2hpi. Boxes outline the 25-75 percentiles, horizontal black lines denote the median. **d-e**, UMAP plots colored by normalized expression of representative up-regulated genes in the 2hpi cluster. UMAP projections reconstructed based on the combined mouse and *Plasmodium* transcriptomes. **f**, smFISH images of supposedly up-regulated genes at 2hpi and 24hpi uninfected hepatocyte show no discernible change in expression. Cxcl10 mRNA in red (positive signal in non-parenchymal cell indicated by white arrow on left panel), Mt1 mRNA in green, Dapi in blue.

**Extended Data Fig. 3:**
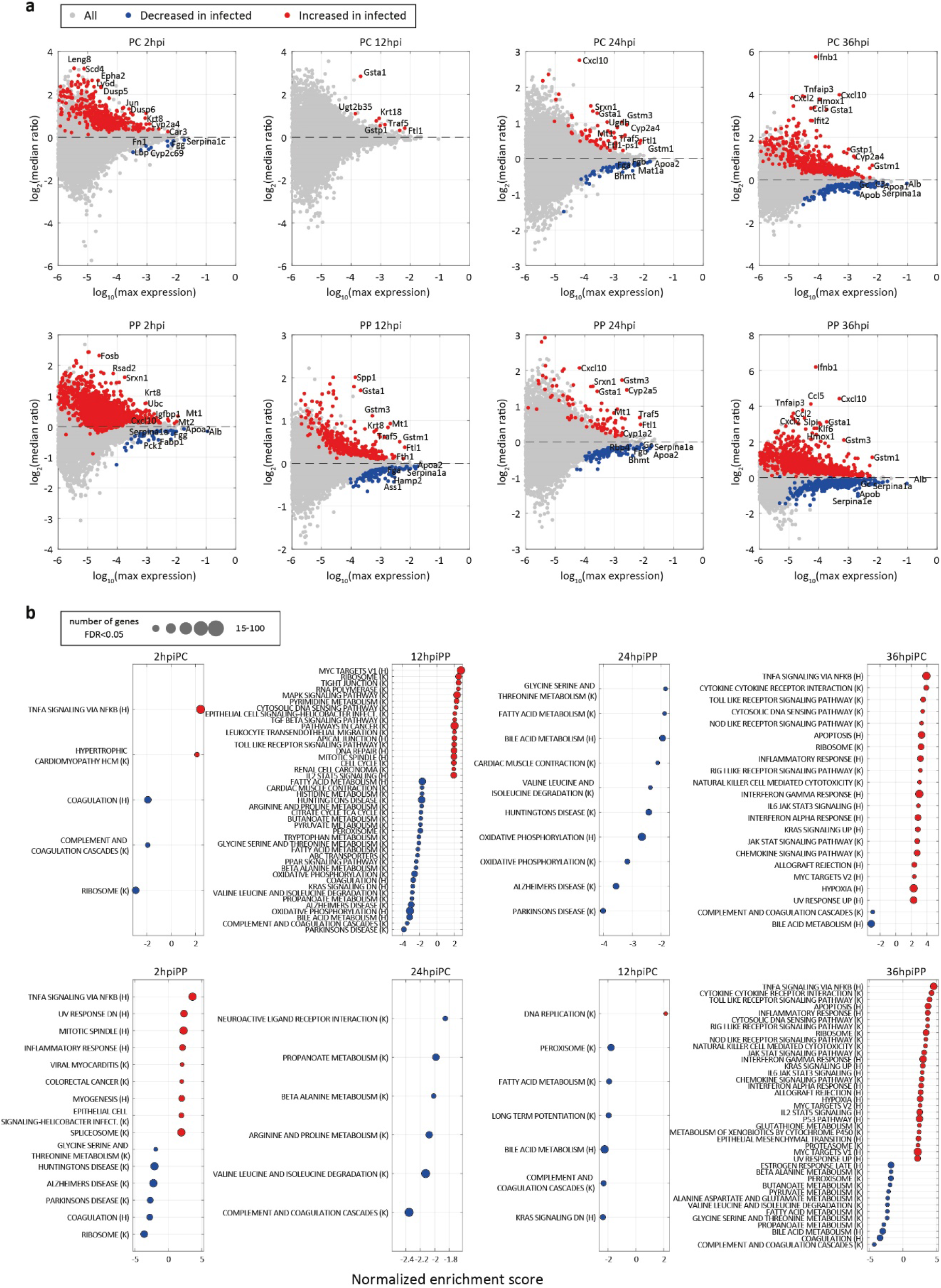
Differential gene expression in different spatio-temporal bins. **a**, MA plots of the median expression ratio between infected and uninfected hepatocytes, split into 8 spatio-temporal bins. Y axis indicates log2 of the median ratio per gene. X axis indicates log10 of the gene’s max expression. Genes significantly increased or decreased in infected hepatocytes are plotted in red or blue respectively (FDR q-value<0.01). **b**, Gene set enrichment analysis (GSEA) for highly expressed genes ranked based on ratio changes indicated in (a). (H) denote Hallmark gene sets, (K) denote KEGG gene sets.

**Extended Data Fig. 4:**
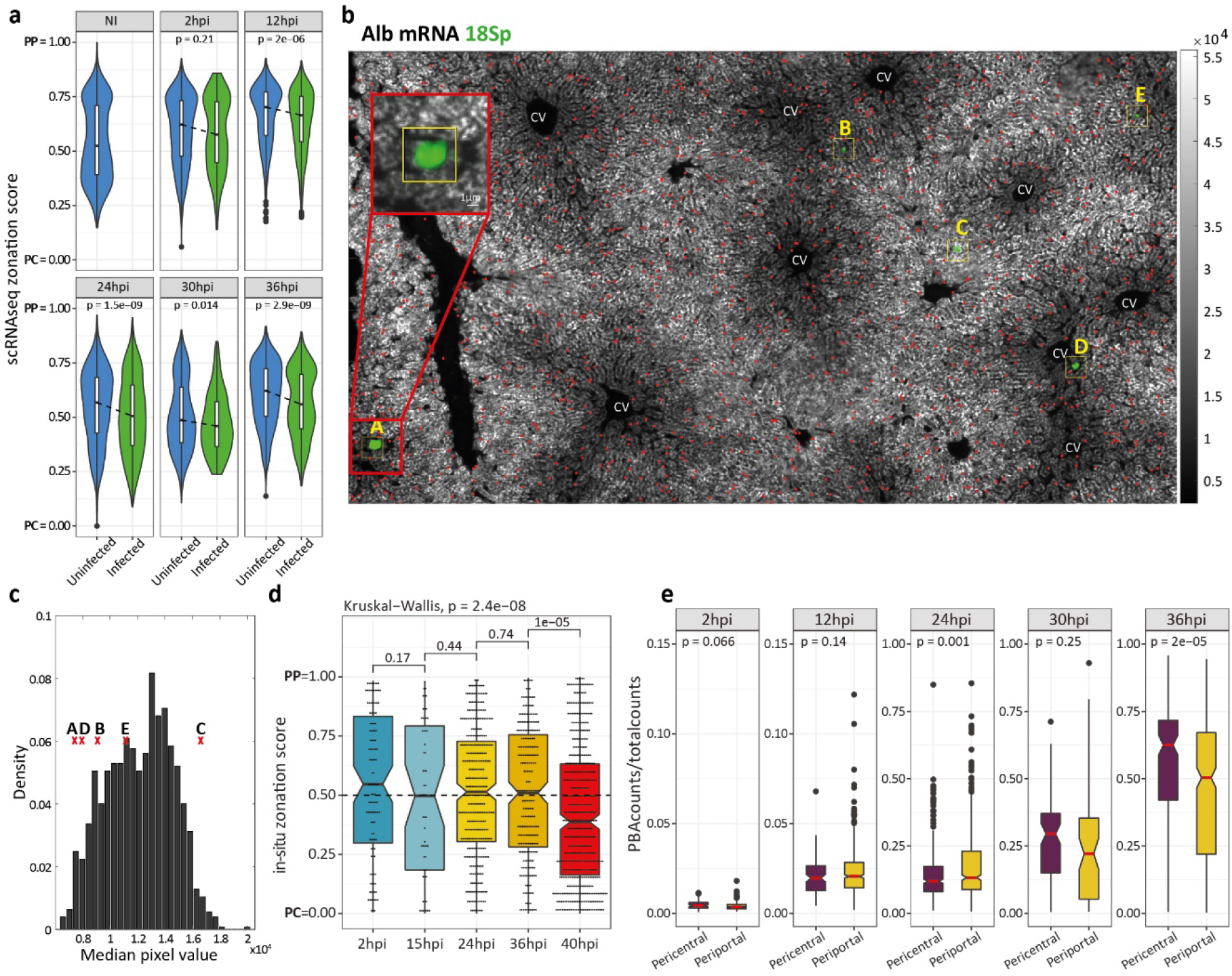
Pericentral infected hepatocytes become more abundant at late time points. **a**, Violin plots of computationally-inferred zonation scores computed on the scRNAseq data. Boxes outline the 25-75 percentiles, horizontal black lines denote the median, dashed line demonstrates the median change between groups. **b**, smFISH of Albumin (Alb, white) and parasite Ch12 18S rRNA (18Sp, green). Yellow squares represents the 40×40 pixels window around the infected cells, number indicates median pixel intensity. Red dots represent the centers of 1248 40×40 pixel windows (“neighborhoods”) randomly placed onto the tissue area. “CV” denotes central-vein locations. Red blow up of 100×100 pixels (11×11 micron) around a pericentral infected cell. Letters indicate the infected cells plotted on **c**, Histogram of median pixel value of random neighborhoods in image, red X represent values of infected neighborhoods labled according to (**b**). **d**, smFISH quantification demonstrates lower proportions of infected periportal hepatocytes at 40hpi. Shown is the zonation score based on the smFISH images, namely the probability to observe a neighborhood zonation lower than the value observed in a random cell within the image field. **e**, scRNAseq analysis demonstrates that periportal parasites have lower transcript counts. For (**d**) and (**e**), boxes outline the 25-75 percentiles, horizontal black/red lines denote the median.

**Extended Data Fig. 5:**
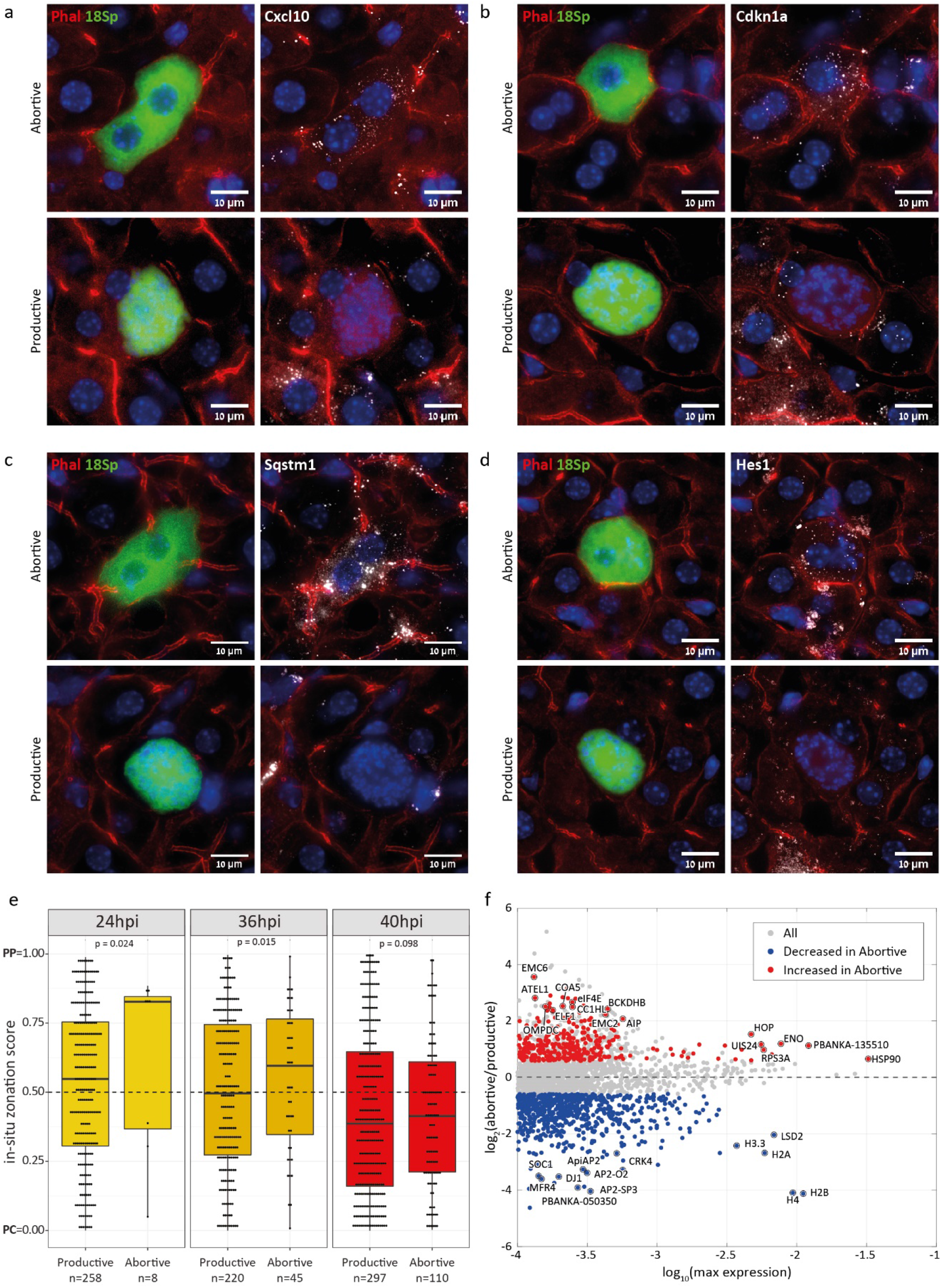
smFISH validations of host genes up-regulated in abortive hepatocytes. **a-d**, Left panel - Phalloidin (Phal) in red, parasite Ch12 18S rRNA (18Sp, PBANKA_1245821) in green, Dapi blue; Right panel – Cxcl10/Cdkn1a/Sqstm1/Hes1 mRNA in white. **e**, Abortive cells are periportally zonated in-situ at 24hpi and 36hpi. Shown is the zonation score based on the smFISH images, namely the probability to observe a neighborhood zonation lower than the value observed in a random cell within the image field, boxes outline the 25-75 percentiles, horizontal black lines denote the median. Significance was calculated using fisher’s method on one-sided Wilcoxon rank-sum test per mouse repeat. **f**, MA plots of the mean expression ratio between abortive and productive hepatocytes at 36hpi. Y axis indicates log2 of the mean ratio per gene. X axis indicates log10 of the gene’s max expression. Genes significantly increased or decreased in infected hepatocytes are plotted in red or blue respectively (FDR q-value<0.01, ratio>1.5).

**Extended Data Fig. 6:**
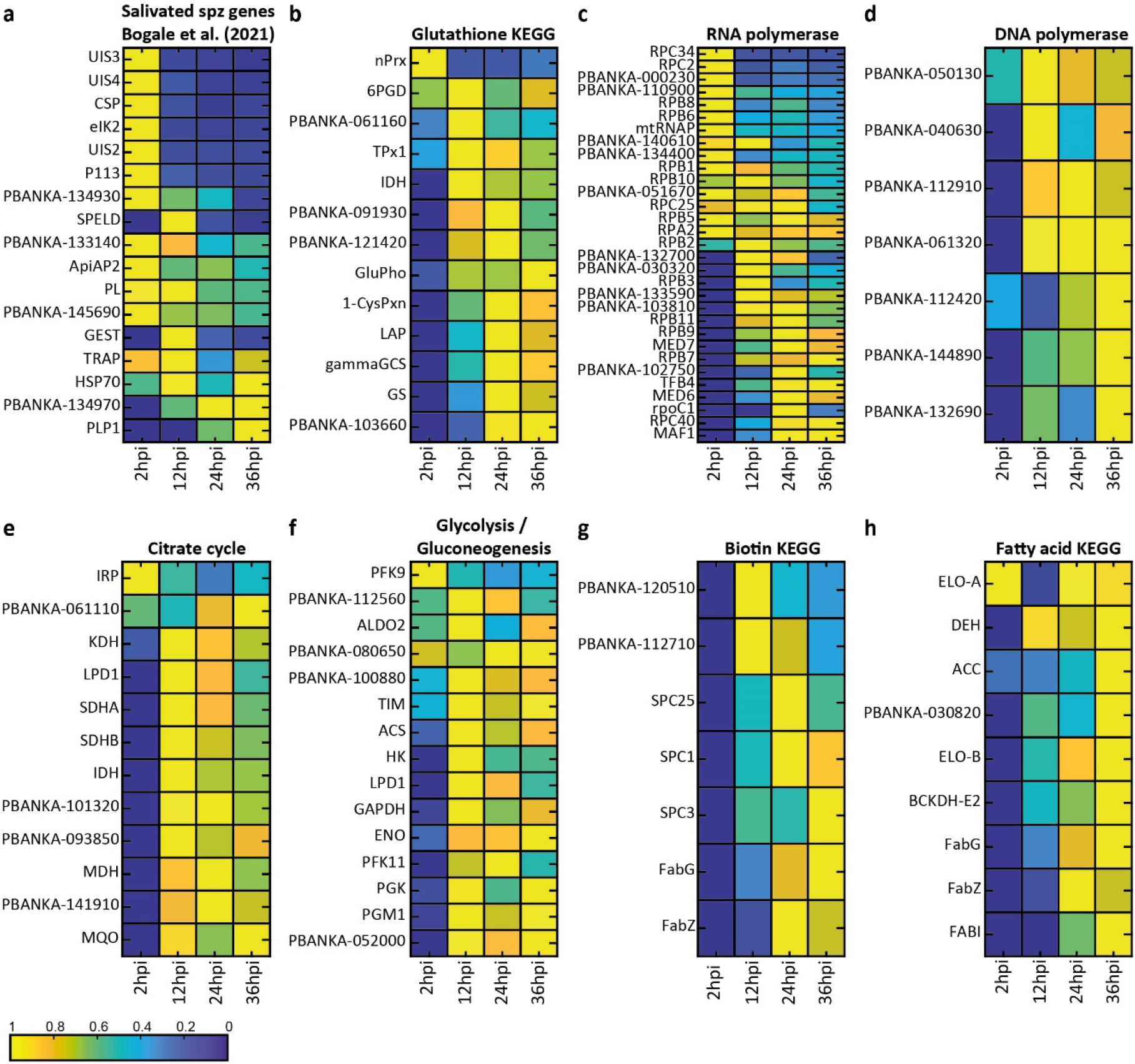
Temporal expression in different *Plasmodium* gene sets. **a**, Marker genes for salivated sporozoites (taken from Bogale et al.^36^). **b**, Genes related to *Plasmodium* KEGG pathway “Gluthathione metabolism”. **c-d**, RNA/DNA polymrases. **e**, Genes related to *Plasmodium* KEGG pathway “TCA cycle”. **f**, Genes related to *Plasmodium* KEGG pathway “Glycolysis / Gluconeogenesis”. **g**, Genes related to Plasmodium KEGG pathway “Biotin metabolism”. **h**, Genes related to various *Plasmodium* KEGG pathways relating to fatty acids (e.g. biosynthesis, elongation, degradation). Color indicates gene-wise normalization over time with yellow=1=max expression.

## Supplementary Tables

**Table S1 - stratified MUS expression over time and space**

Mouse gene expression of zonally-stratified cells at different time points tagged as uninfected (NI) or infected (INF). Shown are means, standard errors of the means (Units are fraction of cellular UMIs), stratified median ratio per gene, one-sided Wilcoxon rank-sum test p-value and Benjamini–Hochberg procedure FDR q-value (left - up regulation in INF, right - down regulation).

**Table S2 - DGE PBANKA**

Differential *Plasmodium* gene expression of (1) zonally-stratified infected cells (2) 36hpi cells tagged as either abortive or productive. Shown are means [Units are fraction of cellular UMIs], ratio between means, two-sided Wilcoxon rank-sum test p-value, and Benjamini–Hochberg procedure FDR (q-value).

**Table S3 - stratified PBANKA expression over time**

*Plasmodium* gene expression of temporally-stratified infected cells. Shown are means, standard errors of the means (se) [Units are fraction of cellular UMIs], center of mass (COM) and KEGG pathways that include this gene.

**Table S4 - Probes used in this study**

Sequences of the smFISH probes libraries used in this study

